# Graph Neural Networks for Likelihood-Free Inference in Diversification Models

**DOI:** 10.1101/2025.08.14.670341

**Authors:** Amélie Leroy, Ismaël Lajaaiti, Sophia Lambert, Jakub Voznica, Maximilian Pichler, Hélène Morlon, Florian Hartig, Laurent Jacob

## Abstract

A common approach to infer the processes that gave rise to past speciation and extinction rates across taxa, space and time is to formulate hypotheses in the form of probabilistic diversification models and estimate their parameters from extant phylogenies using Maximum Likelihood or Bayesian inference. A drawback of this approach is that likelihoods can easily become computationally intractable, limiting our ability to extend current diversification models with new hypothesized mechanisms. Neural networks have been proposed as a likelihood-free alternative for parameter inference of stochastic models, but so far there is little experience in using this method for diversification models, and the quality of the results is likely to depend on finding the right network architecture and data representation. As phylogenies are essentially graphs, graph neural networks (GNNs) appear to be the most natural architecture but previous results on their performance are conflicting, with some studies reporting poor accuracy of GNNs in practice. Here, we show that this underperformance was likely caused by optimization issues and inappropriate pooling operations that flatten the information along the phylogeny and make it harder to extract relevant information about the diversification parameters. When equipped with PhyloPool, a new time-informed pooling procedure, GNNs show similar or better performance compared to all other architectures and data representations (including Maximum Likelihood Estimation) that we tested for two common diversification models, the Constant Rate Birth-Death and the Binary State Speciation and Extinction. We conclude that GNNs could serve as a generic tool for estimating diversification parameters of complex diversification models with intractable likelihoods.

## Introduction

Species richness varies greatly across taxonomic groups (G. E. Hutchison 1959), geological times (Barnosky et al. 2011) and geographical regions (Gaston and Blackburn 2000). It is generally accepted that these patterns in species richness emerge from variation in diversification rates (i.e. speciation and extinction rates) in addition to dispersal, time of divergence and migration processes. For instance, important ecological phenomena such as the ‘Latitudinal Diversity Gradient’ (Hillebrand 2004) may be partly explained by variations in species net diversification rates defined as the imbalance of the speciation and the extinction rate (Mittelbach et al. 2007; Rolland et al. 2014; Pontarp et al. 2019). Our general understanding of the processes that cause these rate changes, however, is still very limited (Rabosky 2009a, 2009b; Condamine et al. 2013; Moen and Morlon 2014).

To better understand the drivers of variation in diversification rates and their consequences for biodiversity dynamics, a common approach is to first accurately estimate net diversification rates from reconstructed phylogenies (Pyron and Burbrink 2013), and then decompose them into speciation and extinction rates (Stadler 2013). The latter is not trivial, as reconstructed phylogenies do not include information on extinct species unless fossil data are available. Without further constraints, the problem is ill-posed, meaning that different diversification dynamics can lead to the same extant phylogeny (Louca and Pennell 2020; Morlon et al. 2022). However, when making additional assumptions about the functional form of extinction and speciation rates over time, it is often possible to statistically estimate the parameters of these functions, and thus infer a probable diversification history that led to the currently observed species richness (Nee et al. 1994; Etienne and Rosindell 2012; Morlon 2024).

One of the simplest diversification models of this kind is the Constant Rate Birth-Death (hereafter ‘CRBD’) model. This model assumes that the speciation and extinction rates are both homogeneous across lineages and constant through time. Under this model, extinction leaves a distinct signal in reconstructed phylogenies called the “pull of the present”, and both extinction and speciation rates can be estimated with Maximum Likelihood (Nee et al. 1994).

In recent years, many studies have worked on extending this and similar models to account for various types of rate heterogeneity as well as their potential drivers (Morlon et al. 2024). Such models commonly allow for variations of speciation and extinction rates through time (time-dependent models), across lineages (inhomogeneous models) and through dependence on environmental (Condamine et al. 2013) or biotic factors (Etienne et al. 2012). For instance, the time-dependent Birth-Death model (Hallinan 2012) allows to consider homogeneous changes—such as exponential decay—of speciation and extinction rates over time (Rabosky and Lovette 2008; Morlon et al. 2011; Stadler 2011). Other diversification models allow lineage-specific shifts in diversification rates that can be either discrete, as can be expected to occur with the appearance of key innovations (Alfaro et al. 2009), or continuous, as can be expected to occur given the gradual evolution of phenotypes (*e.g.* the Cladogenetic Rate Shift ’ClaDS’ model; Maliet et al. 2019, and the Birth-Death-Diffusion ‘BDD’ model; Quintero et al. 2024). The specific innovations or phenotypes that can drive these shifts can be either implicit, as in the latter models, or explicitly modeled, as in State-dependent Speciation and Extinction (SSE) models. A particular example of this is the Binary State Speciation and Extinction model, hereafter ‘BiSSE’ (Maddison et al. 2007), which considers the effect of a binary state on speciation and extinction rates.

A constant challenge when developing new diversification models is establishing robust methods to fit them to data. For some model structures, the likelihood (*i.e.* the relative probability to observe a reconstructed phylogeny given the model and its parameters) can be computed analytically or approximated numerically (see Nee et al. 1994 for the CRBD model and Maddison et al. 2007 for the BiSSE model). If the likelihood can be computed, model parameters can be inferred using either Maximum Likelihood Estimation (MLE; see for instance Ricklefs 2007) or Bayesian inference (*e.g.* Silvestro et al. 2011). However, likelihoods can quickly become analytically or numerically intractable, limiting our ability to perform inference for any hypothesized stochastic process.

To solve the problem of intractable likelihoods, a number of simulation-based methods, among them Approximate Bayesian Computation (ABC), have been proposed. The ABC approach approximates the posterior distribution of a model by comparing model predictions to data via summary statistics (Csilléry et al. 2010; see Saulnier et al. 2017 for an example). Although ABC can successfully infer parameters from complex diversification models, it critically depends on finding a low-dimensional sufficient set of summary statistics for the approximation to be accurate with a reasonable sample size (Hartig et al. 2011). In practice, these requirements have been difficult to achieve, in particular for parameter-rich models.

Recent advances in the field of deep learning provide an alternative approach to likelihood-free, simulation-based inference for phylogenetic diversification models (Lueckmann et al. 2021; Cranmer et al. 2020). In particular, neural estimation, a specific approach to simulation-based inference, trains a neural network on simulations from a probabilistic model to perform inference on its parameters (Lueckmann et al. 2021). In contrast to ABC and related methods, neural estimation relies on the neural network to extract statistics from the data that are relevant to parameter estimation, thus eliminating the need to manually select appropriate summary statistics.

It was shown that neural estimation based on Convolutional Neural Networks (CNN) can outperform ABC for inference in population genomics (Chan et al. 2018; Schrider and Kern 2018). There is, however, little experience on neural estimation in phylogenetic diversification models, and there are many options to implement a neural estimation pipeline. Any neural network (i.e., any architecture with any particular choice of network weights) acting on phylogenies can be thought of as mapping a phylogeny to a set of descriptive statistics (the output of the network). As a consequence, each choice of a particular neural architecture corresponds to a family of descriptive statistics for an observed phylogeny, parameterized by the network weights, and training the network can be thought of as selecting one mapping from phylogenies to descriptive statistics within this family.

When setting up a neural estimation procedure, an obvious requirement is that the neural network must be able to process a phylogeny as input, and predict parameters as output. This is not entirely trivial, as a phylogeny is a relatively complex data structure that cannot be directly processed by any machine learning algorithm. The simplest solution is to summarize the shape of the phylogeny by a suitable set of summary statistics, which allows using practically all common machine learning algorithms that work on tabular data for the neural estimation. The drawback of this approach is that, as for ABC, it relies on finding appropriate (although not necessarily low-dimensional) summary statistics that extract all relevant information from a phylogeny, which is hard to guarantee.

The next option is to transfer the full information contained in a phylogeny into a data-structure that can readily be used by standard deep learning approaches. The most obvious choice here is a string or regular grid. Voznica et al. (2022), for example, developed a bijective matrix representation for non-ultrametric phylogenies with the goal of fitting epidemiological models. They compared the performance of likelihood-based inference techniques to neural estimation using a classic multilayer perceptron (MLP), which consists of fully connected layers with a nonlinear activation function, and to neural estimation using CNN that were supplied either with this matrix representation or with conventional summary statistics. Lambert et al. (2022) adapted this approach for fitting birth-death diversification models to reconstructed (ultrametric) phylogenies, extended the matrix representation to the case of representing phylogenies with associated tip state data in the form of “Complete Diversity-reordered Vector” (CDV), and compared the performance of MLP, CNN and MLE. These studies showed that the matrix encoding processed by CNN performed very well, leading to good performance compared to likelihood-based methods (Voznica et al. 2022, Lambert et al. 2022).

Encoding the phylogeny as a matrix, however, “hides” the neighborhood relationships between nodes (i.e., they are not directly given to the neural network), and these relationships could provide important information. They are preserved if the phylogeny is instead simply represented in its original form as a graph, which can be processed by Graph Neural Networks (GNNs). GNNs generalize the idea of CNNs, which require Euclidean neighborhoods, to graphs (Scarselli et al. 2009; Zhang et al. 2019; Wu et al. 2021). Their key advantage over CNNs is that they explicitly use the graph topology to build vector representations – known as embeddings – for each node. These embeddings are iteratively updated by integrating the information from neighboring nodes in the graph. Thus, in principle, using GNNs should be superior to the matrix representation proposed by Voznica et al. (2022). However, previous attempts to use GNNs for phylodynamic inferences in diversification and epidemiological models (Sun et al. 2024; Qin et al. 2024; Perez et al. 2024), including a previous version of this manuscript (Lajaaiti et al. 2023), have provided contrasting results.

In this study, we introduce PhyloPool, a time-aware pooling procedure for graph neural networks. We conjecture that the previously reported contrasting performances of GNNs for phylogenetic neural estimation may stem in part from the use of standard global pooling operators, which are commonly applied in the final step of GNN architectures. Global pooling merges the embeddings of all nodes into a single vector through averaging. While beneficial in other applications, the global averaging could erase critical macroscopic information on the shape of the phylogeny. In particular, it is well known that, Lineage-Through-Time (LTT) plots contain essential information in their initial and final slopes, which are key to estimate parameters of the CRBD model (Nee et al. 1994). This localized information may be diluted or lost when processed through global pooling layers. To solve this problem, PhyloPool performs a pooling that preserves this critical temporal information.

To assess the advantages of the GNN architecture, we compare neural estimation for the likelihood-free parameter estimation in diversification models with different choices of neural architectures, corresponding to different data representations for the phylogeny (see Fig. 1), including: i) an MLP over hand-designed summary statistics (pink in the Fig. 1), ii) CNN over the LTT or CDV representation (green and orange in the Fig. 1), and iii) graph convolutions followed by either PhyloPool or a global pooling layer (blue in the Fig. 1). We show in particular that neural estimation using graph convolutions followed by PhyloPool achieves similar or superior performance compared to both neural estimation with other architectures and MLE, on data generated under the CRBD and BiSSE diversification models. Our results suggest that GNNs can be used as a generic tool for this task, provided that its architecture is designed to preserve the information relevant to the estimated parameters, and that they are correctly optimized.

**Figure 1.**
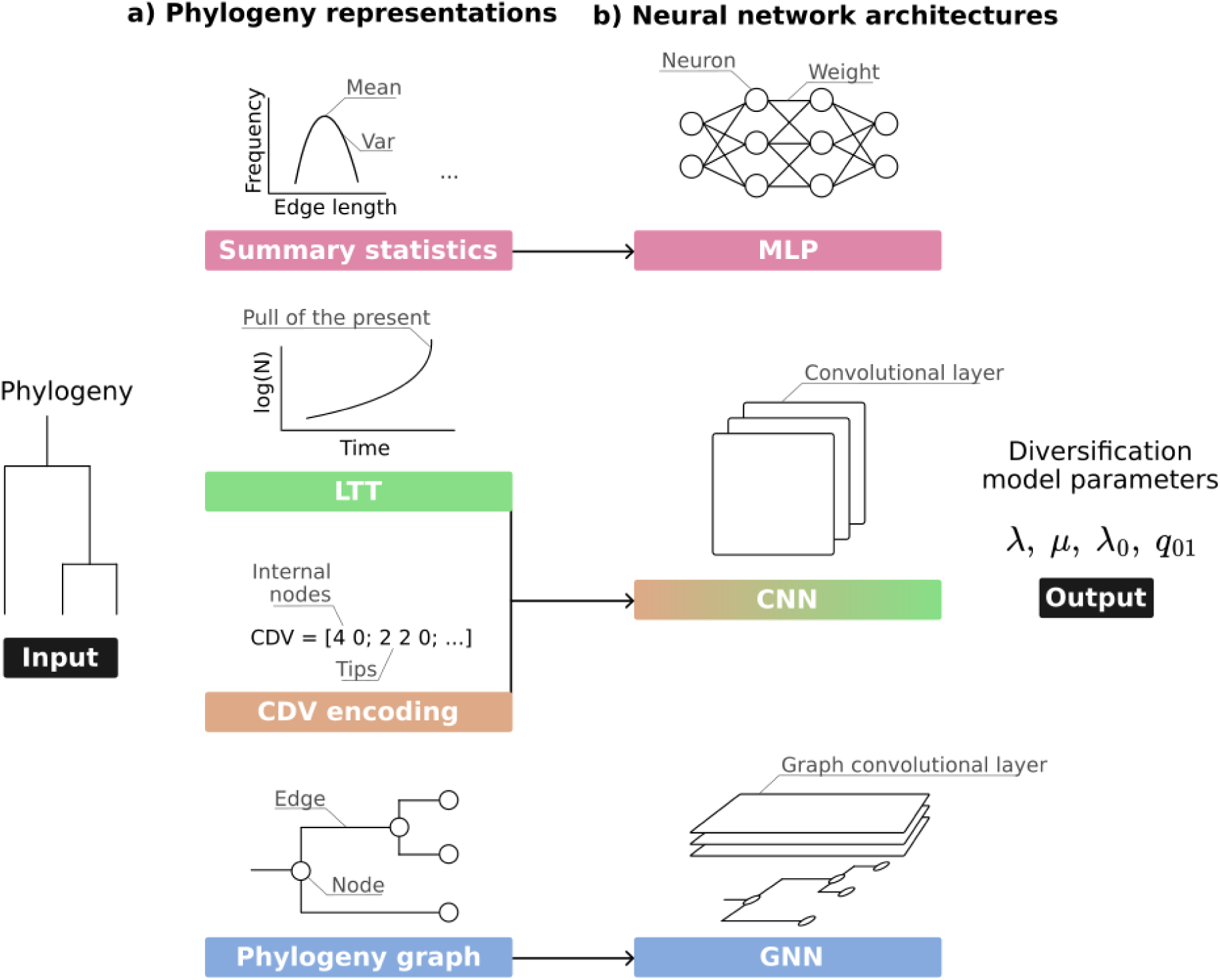
Combining different phylogenetic representations with corresponding neural network architectures. The four considered combinations of phylogenetic representations and neural network architectures are indicated by arrows from column a) to column b): 1) Summary statistics with MLP, 2) CDV with CNN, 3) LTT with CNN, 4) Phylogeny graphs with GNN. The last column describes the output of the inference methods for the two diversification models examined (CRBD and BiSSE).

## Methods

### Diversification Models

We chose two established diversification models as case studies to test the performance of neural estimation for parameter inference in diversification models: 1) the homogeneous CRBD model, and 2) the more complex inhomogeneous BiSSE model. The rationale for choosing these is to have two models with tractable likelihoods but different complexity. In particular, while the CRBD is a homogenous birth-death model, meaning that the LTT contains all information needed for inference with this model, the BiSSE model is an inhomogenous birth-death model with rates that vary across lineages, making the LTT an insufficient statistics for inference.

The Constant Rate Birth-Death (CRBD) model is one of the simplest diversification models. It has two parameters, the speciation rate (λ) and the extinction rate (μ), that are homogeneous across lineages and constant over time. The likelihood of this model is well-known (Nee et al. 1994) and depends only on the phylogeny branching times and not on its topology. To estimate diversification rates with the MLE, we used the APE R package (Paradis et al. 2004).

In the Binary State Speciation and Extinction (BiSSE) model, lineages transition between two states (0 and 1), where each state has its own speciation and extinction rate. Transitions from one state to another over time represent changes in character states that impact diversification rates, such as a shift from sexual to asexual reproduction. The model has 6 parameters: 2 speciation rates (λ_0_, λ_1_), 2 extinction rates (μ_0_, μ_1_) and 2 two transition rates (q_01_ for transition 0 → 1; q_10_ for transition 1 →0). As for the CRBD model, the likelihood of the BiSSE model can be computed (see Maddison et al. 2007). We imposed four constraints on the model to simplify inference, thus reducing the number of free parameters from six to two. The constraints are as follows: 1) λ_1_ = 2λ_0_; 2) μ_0_ = 0; 3) μ_1_ = 0; 4) q_01_ = q_10_. Constraint 1) ensures that states 0 and 1 have different speciation rates, 2) and 3) make the model pure birth, and 4) is an assumption of symmetry that makes the probabilities to switch from one state to another equal. Lambert et al. (2022) also used the CRBD model, and a less constrained version of BiSSE, and showed that their parameters can be inferred using CNN on the CDV representation. Here, we simplified the model to test several network architectures and data representations while limiting the number of simulations (and therefore the computational cost) required to properly train each of the networks. To estimate diversification rates with the MLE, we used the Diversitree R package (FitzJohn 2012) and constrained the likelihood with the 4 constraints listed above.

To train the neural networks, we simulated 100,000 phylogenies for the CRBD and 1,000,000 for the BiSSE model using the Diversitree library (FitzJohn 2012) in R (R Core Team 2023). We assumed complete sampling of the phylogenies, that is no missing species. The latter assumption could be relaxed, as in (Lambert et al. 2022), but again we simplified the model to reduce computational cost. For the tree size, we sampled the number of tips uniformly between 100 and 1,000 in all simulations to ensure that models learn to infer model parameters independently of tree size. For the CRBD model, we draw the underlying parameters as follows: 1) we draw uniformly λ in [0.1,1.0]; 2) we draw uniformly the turnover rate ε in [0, 0.9] from which we compute the extinction rate μ = ελ ∊ [0, 0.9λ]. By doing so, we avoid the critical case where λ ≲ μ (*i.e.,* speciation rate is inferior to or equivalent to the extinction rate). For the BiSSE model, we took λ_0_ ∊ [0.1,1.0] to stay in the same range as for the CRBD model and q_01_ ∊ [0.01,0.1] to ensure that one state is not overly represented compared to the other one.

### Neural Estimation

Neural estimation (sensu Lueckmann et al. 2021) is used to estimate model parameters without requiring an analytic formula for the likelihood P(phylogeny∣diversification parameters). The approach consists of generating data samples (in our case the phylogeny), with their corresponding diversification parameters, and then use these parameter-data pairs to train a neural network with the goal of predicting the model parameters from a given phylogeny.

It can be shown that if i) the model parameters are drawn from a certain prior distribution P(diversification parameters), ii) phylogenies are simulated from P(phylogeny∣diversification parameters) using a probabilistic diversification model, such as CRBD or BiSSE, and iii) the loss function used to train the network is the MAE (see *Training Neural Networks* section below), the trained network will provide a point estimate that converges (for many simulations) to the median of the posterior P(diversification parameters|phylogeny) for the likelihood of the stochastic process (Shynk 2012, 9.12). This statement assumes that the neural network is flexible enough to produce unbiased predictions and that we manage to minimize the MAE through numerical optimization. In the Discussion, we extensively discuss how these three possible limitations (insufficient training data, network expressivity or numerical optimization) affect the accuracy of our approximation for the posterior median. Note that it would also be possible to extend the approach to estimate the full posterior distribution through neural estimation, e.g. by using mixtures of Gaussians, or normalizing flows (Radev et al. 2022), which could be a goal for further research.

### Phylogeny Representation

The success of the neural estimation procedure depends on the ability of the neural network to build a representation of the data that contains sufficient information on the parameter to be estimated. This representation will in general depend both on the input provided to the neural network and on the operations performed by the network over this input. Here we describe the combinations of input and network architecture choices that we consider in this study. We provide a more detailed description of all architectures in the Appendix.

*Summary statistics + MLP (represented in pink in Fig. 1) —* Arguably the most basic option is to represent the phylogeny by a number of summary statistics. We used a set of 84 summary statistics inspired from the set of (Saulnier et al. 2017). Those summary statistics can be split into three groups:

1. 8 statistics related to phylogeny topology (*e.g.* ratio of the width over the depth of the phylogeny);
2. 25 branch lengths statistics (*e.g.* median of all branch lengths);
3. 51 LTT statistics (binned LTT coordinates and LTT slopes).

The list of the summary statistics and the changes compared to (Saulnier et al. 2017) are detailed in the Table S0 of the Appendix. For the BiSSE model we use the ratio of the number of tips in state 1 over the total number of tips as an additional summary statistic. We recognize that more informative summary statistics could be designed, in particular statistics combining the state information with associated phylogenetic branch-lengths, or measures of phylogenetic signal, but avoiding the need to fine-tune summary statistics is one of the main goals of developing deep learning approaches. The resulting set of summary statistics has no particular structure (e.g. graph or sequential) and we pass it through a multilayer perceptron (MLP) to predict the parameters of interest (see Table S1 in the Appendix for details on the MLP).

*Lineage Through Time (LTT) + CNN (represented in green in Fig. 1)—* LTTs illustrate the increase in lineages over time. For a reconstructed phylogeny of n tips, the LTT is composed of n-1 points where each point is defined by two coordinates: time (t, abscissa) and the number of lineages (N, ordinate). For a binary tree where each branching event results in two daughter lineages, the LTT can be compressed to a 1-d array without loss of information, by throwing away the number of lineages and keeping only the times of speciation events, thereby providing a sequence of branching times. The LTT has a sequential structure, we therefore pass it through a convolutional neural network (CNN) to predict the parameters of interest (see Table S2 in the Appendix for details on this CNN).

*Bijective phylogeny encoding (CDV) + CNN (represented in orange in Fig. 1)—* Yet another alternative, already mentioned, is to encode the phylogeny in a regular structure. One option is a real-values vector of length 2n-1 named the ‘Compact Bijective Ladderized Vector’ (see Voznica et al., 2022). Each value of the vector corresponds to one node (internal node or tip), thus there are n values for the n tips and n-1 values for the n-1 internal nodes, which result in a vector of 2n-1 values. The encoding is done in two steps: 1) phylogeny ladderization, and 2) phylogeny traversal. The principle of ladderization is to order each node’s children, such that the encoding is bijective (*i.e.*, one representation encoding maps exactly to one phylogeny and reversely). Here we ladderized the phylogeny such that for each node the left child is the child which is further from the root. On the ladderized phylogeny, we perform an inorder traversal using a classical recursive algorithm. If the visited node is an internal node or the first tip is visited, its distance to the root is added to the vector. Otherwise, its distance from its parent node is added to the vector. Moreover, for the BiSSE model we add the n tip states to the vector. The tip states are ordered according to the phylogeny traversal. Note that in both Voznica et al. (2022) and Lambert et al. (2022), the CDV representation consisted of a two-row matrix, while here we concatenate two 1-d vectors resulting in a 1-d vector. We found that this modification did not affect the results. We passed the CDV representation through a convolutional neural network (CNN) to predict the parameters of interest (see Table S3 in the Appendix for details on this CNN).

*Phylogeny as a graph (GNN) (represented in blue in Fig. 1)—* The disadvantage of encoding the phylogeny in an array-type structure, as in the previous CDV encoding, is that neighborhood relationships of the graph are hidden in the process (i.e., they are not directly given to the neural network). This can be avoided by using Graph Neural Networks (GNN) that process graph-structured representation of the data, such that relationships between nodes in the phylogeny are directly given. In GNNs, each node has an initial embedding vector that is then iteratively updated using the embeddings of its neighbors through graph convolutional layers, a common update scheme. These are followed by pooling layers to capture global features of the graph, and finally fully connected layers. In graph convolutional layers, the update for each node starts from an average of the node itself and its direct neighbors in the graph, normalized by the degree of the nodes (see Fig. 2.a). The new embedding is then formed by applying a linear transformation of this average using a weight matrix learned during training, followed by a pointwise nonlinear function (i.e., applied to each element of the new embedding). After the graph convolutional layers, the output will have a size of *number of nodes × chosen embedding size*. GNNs aiming at predicting global features of the graph (like the diversification parameters in our case) add pooling layers that merge the embeddings of all nodes into a single vector of the same dimension as the embedding size.

**Figure 2.**
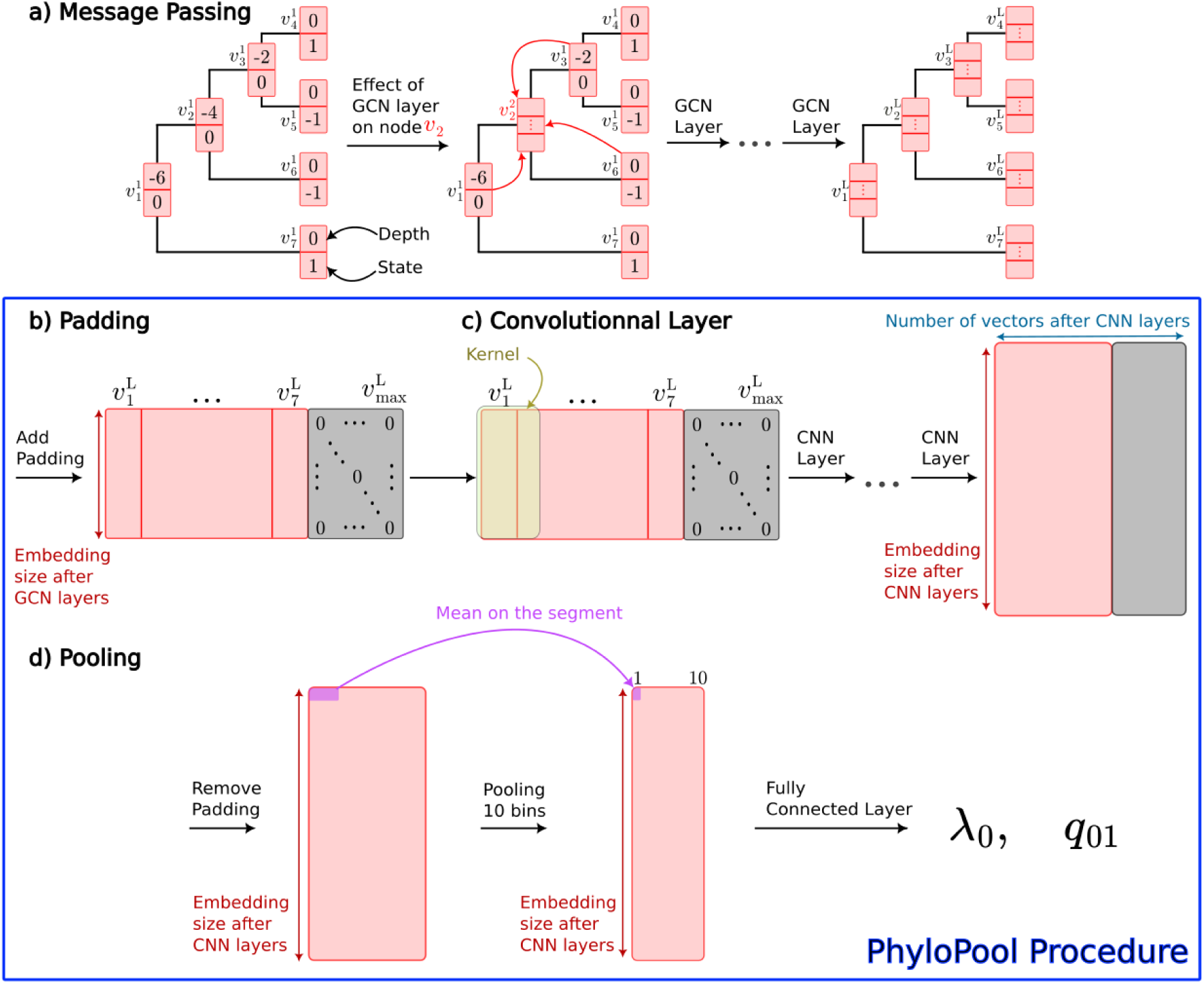
Architecture of PhyloPool for a tree generated under the BiSSE model. a) Message Passing: The first graph represents the initialization: in red, the embedding vector for each node, which includes the depth and state of the node. For example, 𝑣 represents node 2 at the first iteration. In the second graph, we illustrate how a node is updated by incorporating information from its neighbors. This process is repeated over all Graph Convolutional Network (GCN) layers to obtain an embedding of the desired size for each node. b) Padding: We concatenate all these node embeddings into a matrix and add padding up to the maximum (max) number of nodes a tree can have. c) Convolutional Layer: A convolutional layer (CNN) is then applied: a kernel scans across the different nodes, and by repeating CNN, we obtain a new embedding of the chosen size. The number of vectors also changes, as the convolutional layer includes intermediate pooling that reduces the size by a factor of two (see Appendix Table S.5). d) Pooling: Finally, we remove the portion of the convolution output corresponding to zero padding, divide the remaining part into 10 equal segments, compute the mean for each segment, and apply fully connected layers (FC) to these 10 averages. Steps b), c), and d) correspond to the PhyloPool procedure.

A common approach to designing pooling layers is simply to average all embeddings, which is the method used in a previous version of this manuscript (Lajaaiti et al. 2023). In our case, starting from initial node representations that contain the node depth, we know that sorting nodes according to their depth in the phylogeny, which provides a time-series containing the same information as the LTT, should be sufficient to infer the birth and death parameters of the CRBD model. However, averaging across all nodes flattens this information, and coming up with node embeddings that would reproduce the LTT through this flat averaging seems counterproductive (a similar problem is known in other spatio-temporal tasks in the GNN literature, e.g Guo et al. 2019, Su et al. 2020). Instead, we know that the slope of the LTT at the origin of a phylogeny provides a good estimation of the net diversification rate (birth-death), while its slope towards the present provides a good estimation of the birth rate (Nee et al. 1994). This suggests to exploit the ordering of the nodes along the depth of the phylogeny during the pooling step.

To solve this problem, we introduce a time-informed pooling procedure applied after the graph convolutional layers which we call PhyloPool (Fig. 2). PhyloPool applies 1D convolution layers (Lecun et al. 1995) to the matrix obtained by concatenating the node embeddings of the phylogeny, sorted by their depths (Fig. 2.c). To use 1D convolution layers, a fixed input size is required. Because we work with graphs of different sizes, the size of the convolution outcome is determined by the largest graph size allowed, and part of this outcome is thus “padded” (i.e. filled) with zero entries for smaller graphs (Fig. 2.b). Consequently, a small coordinate in this outcome can correspond to information related to nodes relatively close to the root in a large graph or close to the leaves in a small graph. Conversely, a large coordinate can contain information from nodes close to the leaves in a large graph, or zero entries in a small graph. Learning a function that captures critical information such as slopes at the beginning or the end of the LTT from these absolute coordinates using fully connected layers is therefore not optimal. Instead, PhyloPool removes the fraction of the convolution outcome that corresponds to the zero padding, divides the remaining fraction into 10 equal segments, computes the average over each of these segments and applies fully connected layers to these 10 averages (Fig. 2.d). This binning step allows PhyloPool to extract information on relative fractions of the sequence of nodes along the phylogeny in a size-invariant fashion.

We used the Pytorch Geometric framework in Python (Fey and Lenssen 2019) to represent the phylogeny as a graph and train our GNNs. We provide the GNN with the phylogeny’s topology and one attribute per node as its embedding: the distance to the tips, with the tips assigned 0 and the root at a negative coordinate corresponding to the depth of the phylogeny. For the BiSSE model, we add a second attribute containing the node states. 0 is used for tips with unknown states or internal nodes (which state is unknown), and -1 and 1 to encode known states at the tips.

We consider two GNN architectures, both starting with graph convolutional layers. The first architecture (GNN-avg) aggregates the outcome of the last convolution through a global average pooling layer (as in Lajaaiti et al. 2023), the second (GNN-PhyloPool) through our PhyloPool procedure (see Table S4 and S5 in the Appendix for details on both GNNs).

### Training Neural Networks

To train and validate the performance of the networks, we simulated 100,000 phylogenies under the CRBD model and 1,000,000 phylogenies under the BiSSE model. For both models we split phylogenies randomly in three groups: 1) a training set to train the neural networks (90% of the phylogenies); 2) a validation set used to quantify the performance of the neural network during the training (5% of the phylogenies) and 3) a test set also called benchmark data used to quantify the performance of the neural network after the training (5% of the phylogenies). These split sets were the same for all neural network architectures, so all architectures were trained and tested with the same phylogenies.

Training of the networks followed the standard practice of dividing the training data into small batches (typically 64 phylogenies per batch, except for BiSSE with GNN, where we used 128 per batch to reduce the computational time required to go through all batches). To avoid numerical optimization issues, we performed manual hyperparameter tuning for each architecture, guided by the objective that each network should be able to overfit a small dataset. The selected hyperparameters for each architecture are provided in Table S6 in the Appendix. We then trained all networks using the Adam algorithm (Kingma et al. 2017) to minimize the mean absolute error (MAE) for each parameter: 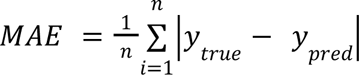, where 𝑦 is the true value for one parameter, and 𝑦 is the value predicted by the network forthat parameter. Since neural estimation approximates the posterior distribution, minimizing the MAE error during training will lead to a point estimate that corresponds to the median of the posterior distribution (in contrast, minimizing the Mean Squared Error loss would have led to the mean of the posterior (Shynk 2012, 9.12)). The mean of the MAEs for both parameters is then used to optimize the network. We used early stopping in the training to avoid overfitting (see Appendix for more details). We found that the value of the weight decay – a parameter used during optimization to prevent the learned weights from becoming too large (equal to L2 regularization) – used in Lajaaiti et al. (2023) was too large in the case of the GNN-avg, and we therefore reduced it to obtain a better optimization (see Appendix for more details).

### Performance assessment

Once the neural networks were trained, we used them to infer the parameter values of the models from the benchmark data (the phylogenies in the hold-out). We also calculated the Maximum Likelihood parameter Estimate (MLE) for these phylogenies to establish a baseline. Thus, for each inference method (the different neural network architectures and the MLE) we have the predicted model parameters (𝑦_𝑝𝑟𝑒𝑑_) and the true values (𝑦_𝑡𝑟𝑢𝑒_) that were used when simulating the phylogeny.

We used two metrics to evaluate the accuracy of the parameter estimates: the MAE, which was the target during neural network training, and the Mean Relative Error (MRE): 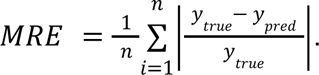. Both metrics frequently appear in simulations-based evaluations of phylogenetic diversification methods (Lambert et al. 2022; Voznica et al. 2022). They are complementary when 𝑦 can take a large range of possible values, since the MAE can remain low while making relatively large errors when 𝑦_true_ is small, while the MRE is very sensitive to large errors on small 𝑦 .

## Results

### Overall Performance of the Architectures

Overall, all methods are able to extract meaningful information from reconstructed phylogenies, although with notable differences (Fig. 3 & 4). In particular, the GNN-PhyloPool architecture performs at least as well, and mostly better than MLE under both the CRBD and BiSSE models, suggesting that we managed to build a sufficient approximation of the posterior with this neural architecture. GNN-PhyloPool also performs at least as well, and mostly better than all other simulation-based alternatives. In contrast, the GNN-avg is the least accurate of all methods for inference of the CRBD model (Fig. 3), and less accurate than CNN-CDV and GNN-PhyloPool for BiSSE (Fig. 4). For inference on CRBD, all neural estimation methods perform at least as well as the MLE, except GNN-avg, and – when evaluating performance using MRE – also except the CNN-CDV for λ estimates, and MLP-SS for μ estimates (Fig. 3). For inference on BiSSE (Fig. 4), CNN-LTT is unsurprisingly the least accurate as no information on tip states is captured in the LTT; MLP-SS, with the restricted summary statistics on tip states provided as input, also exhibits limited performances, comparable to those of GNN-avg; the accuracy of CNN-CDV is quite close to that of MLE, and that of GNN-PhyloPool at least as good, except for when measured with MRE for λ_0_. When they occur, under-performances illustrate possible shortcomings of the neural estimation approximation.

**Figure 3.**
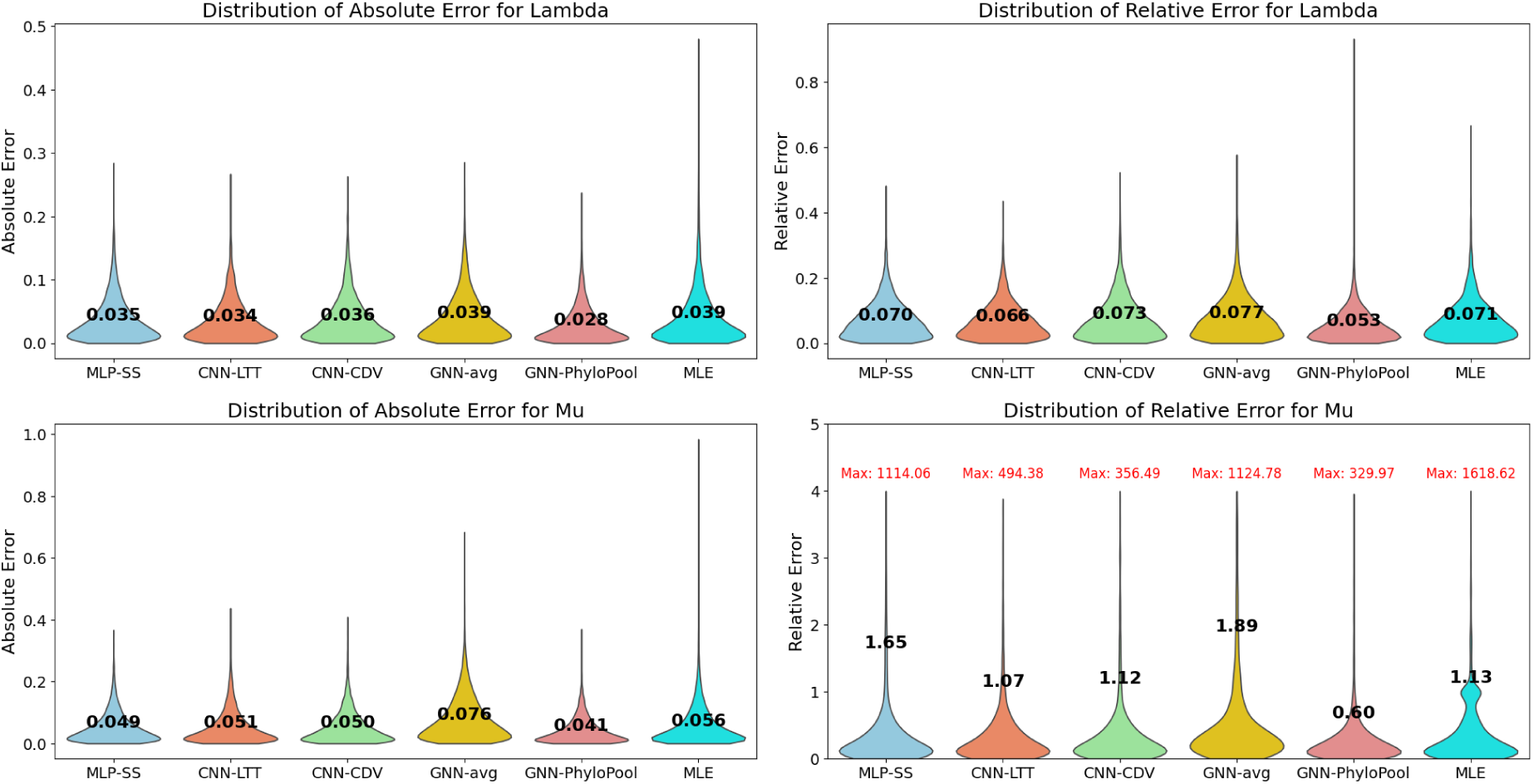
Distribution of Absolute and Relative Errors for the CRBD Model. Distribution of absolute (column 1) and relative (column 2) errors in estimates of speciation (λ) and extinction (μ) rates for machine learning methods and MLE (test set: 5,000 phylogenies). Bold numbers are the means of the MAE and MRE distributions. The machine learning methods evaluated are the following: MLP with summary statistics (MLP-SS), CNN with LTTs (CNN-LTT), CNN with CDV representation of phylogenies (CNN-CDV), GNN with phylogeny graphs with the mean pooling (GNN-avg) and GNN with PhyloPool (GNN-PhyloPool). For the Relative Error of Mu, the distribution is not displayed entirely for better visibility; the red "max" value indicates the upper limit of the range. Additionally, we include in the Appendix (Figure 2) scatterplots of the true versus predicted values for each architecture.

**Figure 4.**
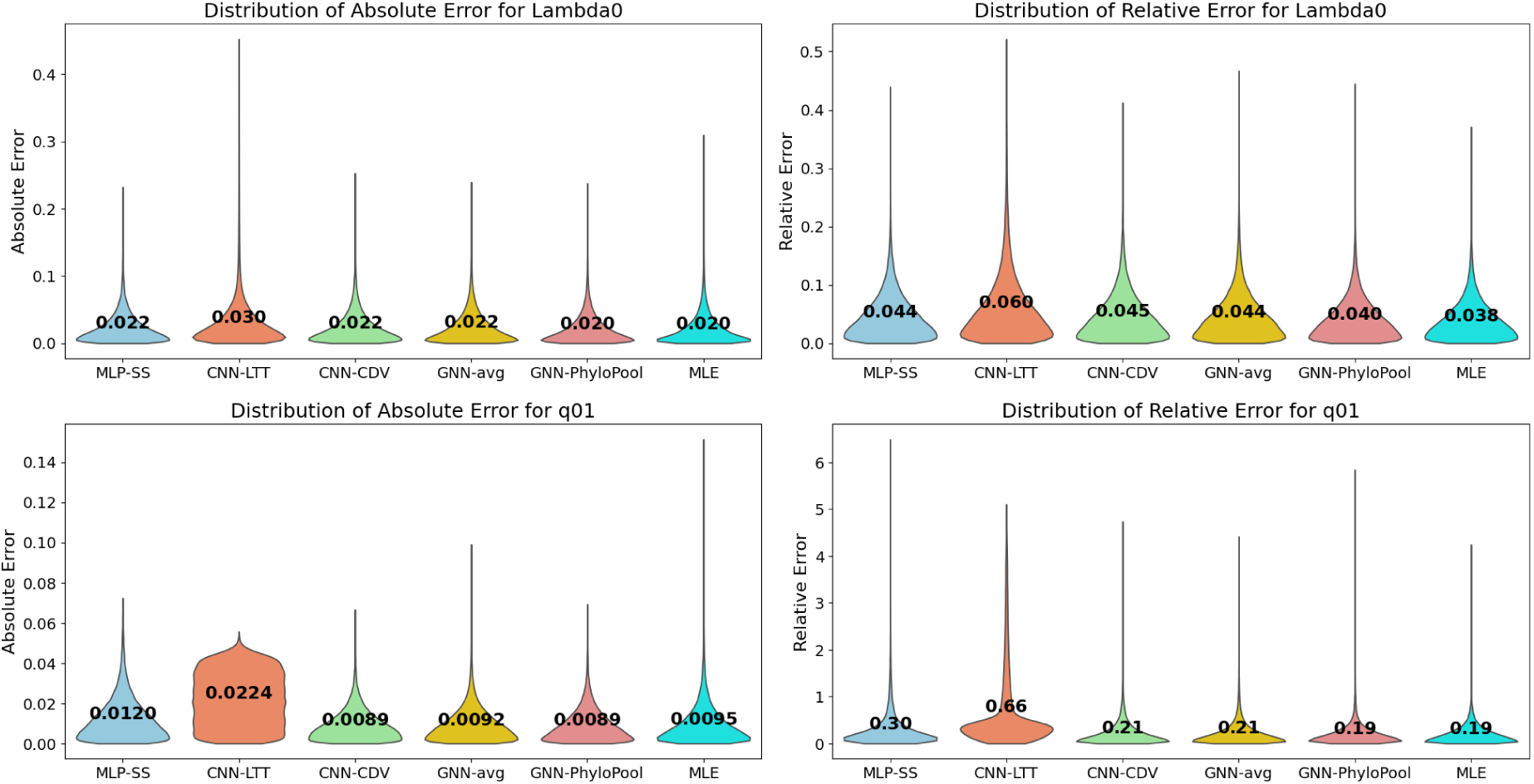
Distribution of Absolute and Relative Errors for the BiSSE Model. Distribution of absolute (column 1) and relative (column 2) errors in estimates of the speciation rate associated with state 0 (λ_0_) and the transition rate (q_01_) for machine learning methods and MLE (test set: 50,000 phylogenies). Bold numbers are the means of the MAE and MRE distributions. The machine learning methods evaluated are as in Fig.3. Additionally, we include in the Appendix (Figure 3) scatterplots of the true versus predicted values for each architecture.

### Importance of capturing the LTT for CRBD

Under the CRBD model and using the MAE metric, all neural estimation methods except GNN-avg performed at least as well as the MLE. This does not come as a surprise for MLP-SS and CNN-LTT which were explicitly designed to capture the slopes of the LTT, which is known to be a sufficient statistic for the parameters of CRBD. GNN-avg on the other hand presumably struggled to recover the LTT because the global pooling step flattens the phylogenetic order of the nodes. This motivated the design of the PhyloPool layer which, when combined with the same graph convolution layers as GNN-avg, led to the best estimate across all methods, suggesting that it succeeded at preserving this information in the LTT while being completely flexible in processing the structure of the phylogeny. We note that optimizing the GNN-avg architecture proved more difficult than for the others architectures: initial attempts with larger values of the weight decay (L2 regularization) hyperparameter – a parameter used during optimization to prevent the learned weights from becoming too large – led to suboptimal solutions where the network predicted a constant value for the death rate μ, as reported in (Lajaaiti et al. 2023). The same difficulty may explain poor performances reported by Perez et al. (2024), whereas Qin et al. (2024) presumably avoid this problem (both references also used a global mean pooling operation as the final pooling step). MLP-SS exhibited intermediate performance in this experiment, likely because it explicitly contains some relevant information on the LTT slopes while being of small dimension and simple to optimize. On the other hand, the fact that it performs less well than GNN-PhyloPool suggests that the 10 local slopes used as summary statistics are not sufficient, and that GNN-PhyloPool manages to capture a little more information from the LTT. More precisely, we hypothesize that the convolutional layers (both over the phylogeny with the graph convolutional layers and along time with PhyloPool) allow each bin to capture useful long-range information compared to slopes computed strictly over 10% windows of the LTT. CNN-LTT and CNN-CDV also obtained intermediate performances, somewhere between GNN-PhyloPool and MLE. We note that, GNN-PhyloPool uses the same CNN architecture as CNN-LTT but differs by its upstream representation, as it starts by graph convolution layers over branching times instead of starting from branching times themselves. These graph convolution layers mix each branching time with its direct neighbors in the phylogeny, which may be far away in the sequence of successive branching times. We hypothesize that under CRBD, this mixing improves the accuracy by allowing the convolution to compute local slopes that are informed and possibly corrected by the rest of the LTT.

Finally, when using the MRE metric, GNN-PhyloPool has an advantage over all other estimation methods, suggesting that other methods sometimes incur relatively large errors on small parameter values, which GNN-PhyloPool manages to avoid, likely because it correctly approximates the posterior distribution and thereby fully exploits the information provided by the prior.

### Importance of capturing the state information for BiSSE

Unsurprisingly on data generated under the BiSSE model, including enough information on the tip states proved to be a decisive factor for good inference. More precisely, MLP-SS, which includes global information on the proportion of the states, was able to accurately estimate the birth rate λ_0_, but not the transition rate q_01_. CNN-LTT, which includes no information on the states, did not provide accurate estimates for either of the two parameters. All other neural estimation methods accounted for tip states and had better performances. Overall, since we need information on topology and tip states (see Fig. 1 in the Appendix), the LTT-based statistics are less useful under BiSSE, which explains why the PhyloPool procedure loses part of its edge against global pooling (used in GNN-avg): preserving the phylogenetic order is intuitively less important when estimating under the BiSSE model, where consecutive nodes may be under different states. However, it remains the best architecture tested, achieving a performance comparable to the MLE.

## Discussion

In this work we introduced a Graph Neural Network with a time-aware pooling layer, PhyloPool, specifically designed to better capture information provided by LTT to estimate the parameters of probabilistic models of diversification. We then compared maximum likelihood estimators to likelihood-free neural estimators exploiting a variety of neural architectures, and showed that the GNN exploiting PhyloPool is at least as accurate, and sometimes better, than all other estimators under the two considered diversification models.

Both MLE and neural architecture obtain parameter estimates under a probabilistic model of the data given the parameters. They differ in the way they access this model: through its likelihood function for MLE, i.e., by evaluating the probability function, or through simulations for neural estimation, i.e., by sampling the probability function. In our experiments, we built our MLE, neural estimation and benchmark data under the same probabilistic models, which excludes misspecification issues. MLE is based on numerical optimization of the likelihood function which is available for both the CRBD and BiSSE models. Neural estimation on the other hand builds an approximation of the posterior distribution through sampling. If the approximation is good enough, it should outperform MLE in our experiments as it exploits a prior distribution which is not available to the MLE (this prior is materialized by the distribution used for sampling parameter values used for training our networks, meaning the prior is given by the simulations.). Since we generate our benchmark data using the same prior distribution, we are concretely reducing the variance of the neural estimates without increasing its bias, which necessarily leads to lower errors. On the contrary, cases of neural estimation leading to larger errors must be explained by the quality of our posterior approximation, i.e., by the inability of the trained neural network to correctly predict diversification parameters for unseen phylogenies, which in turn can be caused by the network building an inefficient representation of the data (e.g., where many statistics are unrelated or indirectly related to the parameter), being trained on too little data, having an insufficient capacity (e.g., too few parameters) or being insufficiently optimized.

Accordingly in learning theory, it is standard to analyse error (sometimes coined “generalization error’) as a combination of three terms denoted approximation, estimation and optimization error (Bottou and Bousquet 2007), linked to inadequate network architecture, insufficiently large training set and optimization issues, respectively. We believe that this analysis could be used as guiding principles when designing neural architectures for neural estimation, and will now illustrate how it sheds a useful light on our results.

A neural architecture defines a family of functions, one for each possible choice of its learnable weights. In the context of neural estimation, the first layers can be thought of as a function extracting a vector of summary statistics from the input phylogeny, and the last layers as a function estimating the diversification parameters from these summary statistics. In other words, designing the architecture amounts to defining a family of possible summary statistic vectors, and training the network amounts to choosing one within the family (by contrast, in ABC, one particular vector of summary statistics is chosen *a priori*). Approximation error is caused by learning with a family of functions (defined through the choice of an architecture) that is too restricted and contains no function making good predictions of the diversification parameters from the phylogeny. In neural estimation we are estimating the posterior distribution, whose only dependency on the data goes through the likelihood function. A large approximation error therefore suggests that none of the functions defined by the chosen architecture extracts sufficient statistics for the diversification parameters. On the contrary, estimation error occurs when learning with a family that does contain functions extracting sufficient statistics but also contains other functions extracting vectors that are not sufficient statistics but that are better than the sufficient statistics on the set of examples (here, phylogenies) used to optimize the weights during training. This happens because the model is trained on a finite set of samples drawn from the prior distribution, rather than on the distribution itself. As a result, the learning process selects one of these functions, leading to poor estimation on new phylogenies (never seen during the training process but simulated under the same model). Finally, optimization error occurs when the numerical training process stops before reaching the function among those defined by the architecture that would give the best prediction on the training data.

For example, the families defined by MLP-SS and CNN-LTT both contain, among many other descriptive statistics, the local slopes of the LTT, which capture most of the relevant information about the birth and death rates under the CRBD model, and therefore incur little approximation error under CRBD. We observed nonetheless that neural estimation exploiting these two architectures performed slightly less well than the one exploiting GNN-PhyloPool. For MLP-SS, this suggests that the chosen local slopes were not entirely sufficient. For the CNN-LTT, universal approximation theorems (Yarotsky 2022) guarantee that there exists a CNN extracting from these branching times a sequence of local slopes of the LTT at any desired granularity, as this sequence is equivariant by translation of the branching times. The universal approximation property states that, given a sufficiently large number of layers and neurons, a neural network can approximate any continuous function on a compact domain. Because the birth and death parameters in a CRBD model are reflected in these slopes, the small gap between CNN-LTT and GNN-PhyloPool is unlikely to arise from the choice of applying convolutions over branching times, but rather from a failure either to choose the right architecture among possible CNNs (e.g. number of layers or dimensions), or to find the right set of weights for this architecture through numerical optimization. On the contrary, the same two architectures (CNN-LTT and MLP-SS) miss information on topology and the leaf traits, making them insufficient (i.e., incurring a large approximation error) under BiSSE. We initially observed poor performance with GNN-avg, which was due to suboptimal hyperparameter tuning. However, after improving the optimization process, GNN-avg provided reasonably accurate estimations. This demonstrates that at least some fraction of the error observed in Lajaaiti et al. (2023) was caused by insufficient optimization (hyperparameter tuning), and not by the insufficient approximation capacity of the GNN-avg architecture.

Estimation error on the other hand happens when training large networks on too few examples. We did not encounter this situation because we focused on networks with a reasonably small number of parameters and, in the context of neural estimation, we always have the flexibility to simulate additional training examples. A guiding principle for data representation should be the possibility to extract sufficient statistics with simple functions belonging to a restricted class (such as convolutions), which ensures both a low approximation and estimation error.

This observation sheds light on the more general difficulty to design versatile architectures for neural estimation. Graph neural networks are good candidates but led to disappointing performances in previous attempts by us (Lajaaiti et al. 2023) and others (Perez et al. 2024). Our analysis suggests that these results arose from a combination of optimization and approximation errors: once correctly optimized, our GNN-avg led to intermediate performances under CRBD and performed almost as well as GNN-PhyloPool under BiSSE. To minimize optimization error, we recommend that practitioners verify the ability of their network to reach an arbitrarily low value for the loss function over a small training dataset. It serves as a diagnostic step: if the network struggles to minimize the loss in this setting, it may indicate optimization problems resulting from a poor choice of hyperparameters. This step is independent of the model’s final generalization performance but helps ensure that the optimization process itself is functioning properly before applying the network to neural estimation.

Our newly introduced pooling operator PhyloPool further improved the performances of the GNN under both diversification models, likely by reducing the approximation error incurred by the average pooling layer. The design of PhyloPool was guided by our understanding of the sufficient statistics for CRBD, and the principle of building families of functions that are small but contain good approximations. By contrast, pooling all node embeddings through a flat average can probably reproduce an LTT, but likely at the cost of building more complex node embedding upstream, which in turns requires larger networks, more data and a more difficult learning procedure.

Admittedly, PhyloPool may not be appropriate for all probabilistic models of diversification: it was motivated by CRBD and already loses some edge under BiSSE compared to other architectures (although still retaining the best performance). Specifically, it may be less suitable for birth-death models including branch heterogeneity, where the global order provided by the node depths (LTT) becomes a less relevant information. We anticipate that layers that account for this depth order, e.g. through convolutions or possibly self-attention (as used in spatio-temporal graphs (e.g. Guo et al. 2019, Su et al. 2020)), will often be complementary to other layers acting on the topology (encoded in the phylogenetic graph), e.g. through graph convolutions. Common strategies to combine different notions of neighborhoods (like depth and topology) include alternating between the corresponding types of layers or integrating the neighborhood information in the node embeddings as in graph transformers (Rampášek et al. 2022). In addition, graph transformers are known to avoid the common oversquashing issue of other graph neural networks (Topping et al. 2022), where information fails to flow between distant nodes on the graph. This phenomenon is particularly acute on poorly connected graphs such as trees, making graph transformers good candidates for neural estimation on phylogenies that will be worth testing in the future. Their main known caveat is that they typically require more training data than other architectures to show their potential, but this is not necessarily an issue in the neural estimation framework where we are only limited by the computational cost to simulate and train with more data.

Like all Bayesian inference methods, the performance of neural estimation is affected by the relevance of its prior, and real word estimation problems sometimes require very diffuse priors for some parameters. In neural estimation, contrary to likelihood-based Bayesian inference, diffuse priors raise the additional issue to sample sufficient training data to ensure a correct approximation of the posterior over their entire support. Future work should therefore assess the empirical ability of neural estimation to produce accurate estimates of the posterior under diffuse priors. Finally, current neural estimation methods for diversification parameters have been restricted to architectures providing point estimates, typically approximating the median or mean of the posterior distribution. Other choices of architectures and loss functions could provide a more comprehensive quantification of the uncertainty through full posterior distributions (Radev et al. 2022). We are confident that exploring these potential improvements of simulation-based inference, and in particular exploiting the power of GNNs to analyze data that has a natural graph structure – such as the phylogenetic trees of species – will open the door to estimating diversification parameters under more and more realistic scenarios, such as those provided by mechanistic eco-evolutionary models that are currently not amenable to inference (Hagen et al. 2021). Working under these realistic scenarios in turn should provide finer results from empirical phylogenies where common simplifying assumptions may affect current estimates of diversification.

## Supplementary Materials

The code to reproduce the results in this study is available at https://github.com/ameliemelo/Phylo_Inference. Simulation data and the online appendix are available https://zenodo.org/records/16794902?preview=1&token=eyJhbGciOiJIUzUxMiJ9.eyJpZCI6IjRiODU4MGQ5LWNlYWMtNDA0Ni04MDY0LWJmZWYxMWIwZWEyMyIsImRhdGEiOnt9LCJyYW5kb20iOiJhYjljNzkzNTgyOGNhMWU2NGZiYTA3M2MzZGY4NmM4NSJ9.agigUYbp-JWjLSoXmwKY3wmrquj5n1dC1TerdJf5pSz2JKw_HcPpNNJtlC0C2_1GQ7TfnrTeb34EsKf-H9bFtQ.

## Acknowledgements

We thank Sonia Kéfi for helpful discussions and comments on the manuscript. The design of our study profited from the results of an earlier MSc Thesis by Felix Gottschlich at the University of Regensburg that examined the possibility to use GNNs for phylogenetic diversification inference.

Data Availability Statement

